# Cryopreservation impairs cytotoxicity and migration of NK cells in 3-D tissue: Implications for cancer immunotherapy

**DOI:** 10.1101/812172

**Authors:** Christoph Mark, Tina Czerwinski, Susanne Roessner, Astrid Mainka, Franziska Hörsch, Lucas Heublein, Alexander Winterl, Sebastian Sanokowski, Sebastian Richter, Nina Bauer, Gerold Schuler, Ben Fabry, Caroline J. Voskens

## Abstract

Natural killer (NK) cells are important effector cells in the immune response to cancer. Clinical trials on adoptively transferred NK cells in patients with solid tumors, however, have thus far been unsuccessful. As NK cells need to pass stringent safety evaluation for clinical use, the cells are cryopreserved to bridge the necessary evaluation time. While a degranulation assay confirms the ability of cryopreserved NK cells to kill target cells, we find a significant decrease of cytotoxicity after cryopreservation in a chromium release assay. We complement these standard assays with measurements of NK cell motility and cytotoxicity in 3-dimensional (3-D) collagen gels that serve as a substitute for connective tissue. We find a 5.6 fold decrease of cytotoxicity after cryopreservation and establish that this is mainly caused by a 6-fold decrease in the fraction of motile NK cells. These findings may explain the persistent failure of NK cell therapy in patients with solid tumors and highlight the crucial role of a 3-D environment for testing NK cell function.

**Synopsis:** Cryopreservation of natural killer (NK) cells dramatically impairs their motility and cytotoxicity in tissue. This finding may explain the persistent failure of clinical trials in which NK cell therapy is used for treating solid tumors.

## Introduction

Natural killer (NK) cells are important effector cells in the early innate immune response to various pathogens, including cancer. For the elimination of target cells, NK cells form a temporary immune synapse with the target cell and secrete granules containing cell-toxic granzymes and perforin. To reach target cells outside the blood stream, NK cells can extravasate and migrate through the connective tissue of numerous organs (1). Adoptive transfer of human NK cells in mice has been shown to suppress the development of primary tumors and metastases (2–4), and clinical studies have shown encouraging results in patients with hematological malignancies (5,6). Studies in patients with solid tumors, however, have thus far failed to demonstrate antitumor responses (7–11), and although the transferred NK cells remain viable in the peripheral circulation, they seem to lose their cytotoxic function in vivo (10).

The cytotoxicity of NK cells is typically evaluated in CD107a degranulation and chromium-release assays (12–14). The degranulation assay detects CD107a proteins from cytolytic granule membranes that are transported to the surface of NK cells upon formation of an immune synapse. The chromium-release assay measures the actual number of tumor cells that are lysed by NK cells. In both assays, the NK cells are in close contact with the tumor cells and do not need to migrate far to reach tumor cells. In vivo, however, the ability to infiltrate a 3-dimensional (3-D) environment is crucial for NK cells to reach the tumor cells.

Clinical application of NK cells demands stringent evaluation regarding sterility, purity and function. This requires freezing and thawing of the cells to bridge the necessary time for passing pre-defined lot-release criteria. Therefore, clinical studies mainly rely on the use of cryopreserved cells. Cryopreservation has been shown to have no significant effect on NK cell cytotoxicity as measured with classical degranulation and chromium-release assays (15). While our data confirm that NK cells retain their ability to induce target cell death in a degranulation assay, we find a decrease of cytotoxic function in a chromium release assay after cryopreservation.

To investigate the origin of this decreased cytotoxicity, we perform time-lapse imaging of NK cells and target cells embedded in 3-D collagen gels that serve as a model system for the extracellular matrix of connective tissue. We find that the fraction of motile NK cells in 3-D collagen gels is decreased by 6-fold after cryopreservation while the small remaining population of motile cells retains its cytotoxic function in a 3-D environment.

## Methods

### PBMC isolation and storage

NK cells are generated from peripheral blood mononuclear cells (PBMCs) from 5 healthy donors (Department of Transfusion Medicine, University Hospital Erlangen, Germany; IRB approval number 147_13B). Specifically, fresh blood is drawn from the arm vein, from which PBMCs are isolated. PBMCs are re-suspended in human serum albumin (Baxter) at a concentration of 5–10 × 10^6^ cells/ml. 500 µl of cell suspension is filled into a 1 ml cryovial (Nalge, Nunc), and 500 µl of freezing medium (55.5 vol% human serum albumin, 25.0 vol% DMSO, 8 % (w/v) glucose) is added. After gently mixing, the vials are transferred into a freeze container (Mr. Frosty, Thermo Scientific, which allow for a cooling rate of −1°C/minute) and stored at −80°C. Cells are used within 65 days after cryopreservation.

### NK cell expansion

After thawing the PBMC samples (which includes NK, NKT and T cells), cells are expanded in the presence of irradiated K562-mbIL15-41BBL feeder cells (gift from Prof. D. Campana, Department of Pediatrics, University Hospital Singapore) for 14-days in RPMI 1640 medium supplemented with 10% fetal bovine serum, 20µg/ml gentamycin and 1% L-glutamine (hereafter called cRPMI), and 200 IU/ml human IL-2 cytokine (16). K562 feeder cells are confirmed negative for mycoplasma contamination using the Venor GeM Classic detection kit (Minerva Biolabs). This expansion process is performed 3 times in the case of two donors, 2 times in the case of another donor, and one time in the case of the remaining two donors, resulting in a total of 10 expansions.

### Cryopreservation and thawing

Expanded NK cell aliquots (5–10×10^6^ cells/ml) are either directly measured (in the following referred to as “fresh”), or are frozen, thawed, and then measured (in the following referred to as “cryopreserved”). We measured 10 paired (fresh versus cryopreserved) samples in total, but due to technical difficulties in some of the assays, the number of paired experiments varied as noted in the figure legends. Freezing of expanded NK cells is performed as described above for PBMCs. On the following day, cryopreserved expanded NK cells are rapidly thawed in a 37°C water bath until a small visible ice chip remains, and then are drop-wise transferred into 10 ml of cRPMI, which is preheated to 37°C. Cells are centrifuged at 300 g for 5 minutes and resuspended in cRPMI.

### Flow cytometry

Fresh and cryopreserved NK cells are phenotypically characterized as described in (16,17) by staining with directly conjugated mouse anti-human antibodies against CD3 (clone UCHT1), CD56 (clone HCD56) and CD16 (3G8). NK cells are defined as CD3- and CD56+ cells. A minimum of 10,000 cells are analyzed using a BD Canto II flow cytometer (BD Biosciences) and Flowjo Software (FLOWJO, LLC Data analysis software).

### CD107a degranulation assay

1 × 10^6^ expanded NK cells are incubated for 6 hours at 37 °C, 5% CO_2_, 95% RH with cells from the myeloid cell line K562 (gift from Dr. J.J. Bosch, Department of Medicine 5, University Hospital Erlangen) at an NK-to-K562 cell ratio of 20:1 and 5:1 in a final volume of 500μl cRPMI supplemented with anti-CD107a antibody (clone H4A3, 10 µl/ml, BD Biosciences). K562 cells are confirmed negative for mycoplasma contamination. To prevent protein secretion and degradation of internalized CD107a, monensin (1 µM) and brefeldin A (10ng/ml, both from Sigma) are added after one hour of incubation. NK cells alone serve as a negative control, and NK cells stimulated for 6 h with phorbol 12-myristate 13-acetate (PMA, 50 ng/ml) and ionomycine (250 ng/ml, both from Sigma) serve as a positive control for anti-CD107a antibody binding. After 6 hours of incubation, cells are harvested, washed, resuspended in 50μl PBS and stained with live-dead Zombie NIR (BioLegend), anti-CD56 (clone CHD56, BioLegend) and CD16 antibody (clone 3G8, BioLegend). Samples are analyzed using a Becton Dickinson FACS CANTOII flow cytometer and Flowjo software.

### Chromium-release assay

K562 cells are labeled with radioactive (150 μCi, 5.55 MBq) sodium chromate (20 µl/condition, 5 mCi/ml, Perkin Elmer) for 1 hour. After incubation, cells are washed 2 times and incubated for an additional 30 minutes to reduce spontaneous chromium release. Labelled cells are then plated at a density of 5000 cells/well in 100 µl cRPMI in a 96-well U-bottom plate. Fresh expanded or cryopreserved NK cells are added at NK-to-target cell ratios of 1:20, 1:10, 1:5 and 1:2.5 to give a final volume of 200 μl per well. After 4 hours of incubation, 100 μl supernatant is mixed with 100 μl scintillation cocktail (Perkin Elmer) in a 96-well sample plate (Perkin Elmer). Release of radioactive chromium-51 is measured using a gamma-counter (Perkin Elmer), and the fraction of lysed target cells is calculated as the ratio of (experimental release - spontaneous release)/(maximum release - spontaneous release). Spontaneous release is measured from 5,000 labelled K562 cells without addition of NK cells, and maximum release is measured from 5,000 labelled K562 cells that are lysed with 100 µl 1% Nonidet P-40 (Sigma). All experiments are performed in triplicates.

### 3-D cell motility assay

We suspend 150,000 fresh or cryopreserved NK cells in 2.5 ml of a 1.2 mg/ml collagen solution in each well of a tissue-culture treated 6-well plate (Corning). The collagen solution is prepared from a 2:1 mixture of rat tail collagen (Collagen R, 2 mg/ml, Matrix Bioscience) and bovine skin collagen (Collagen G, 4 mg/ml, Matrix Bioscience). We add 10% (vol/vol) sodium bicarbonate (23 mg/ml) and 10% (vol/vol) 10× DMEM (Gibco). For a final collagen concentration of 1.2 mg/ml, we dilute the solution before polymerization with a mixture of 1 volume part NaHCO_3_, 1 part 10 × cRPMI and 8 parts H_2_O (18) and adjust the solution to pH 10 with NaOH.

After polymerization at 37 °C, 5% CO2 and 95% RH for 1 hour, 1.5 ml of RPMI medium is added to each well of a 6-well plate. Following an additional waiting period of 3 hours to ensure that the NK cells have adapted to the collagen gel and have attained their characteristic elongated shape, the 6-well plate is transferred to a motorized microscope (Applied Scientific Instrumentation, USA, equipped with a 10x 0.3 NA objective (Olympus) and an Infinity III CCD camera, Lumenera) that is placed inside an incubator, and time-lapse imaging is started. We perform z-scans (10 µm apart) through the 1mm-thick gel every 30 s for a duration of 5 min. Afterwards, another randomly chosen position is selected, and time-lapse imaging continues. In total, six positions are imaged in each well. Fast image acquisition of z-scans is achieved by taking images at 27.5 frames per second while the focal plane of the microscope is moving with a constant speed through the gel along the z-axis. We do not store the entire z-stack of images but instead store the lowest intensity value of each pixel along the z-direction (minimum intensity projection, Fig. 1a,b) and the z-position where the lowest intensity value is found (Fig. 1c). The z-position information aids in the tracking of two individual cells when their paths seem to cross in the minimum intensity projection images. In a similar way, we determine the maximum intensity projection.

**Figure 1:**
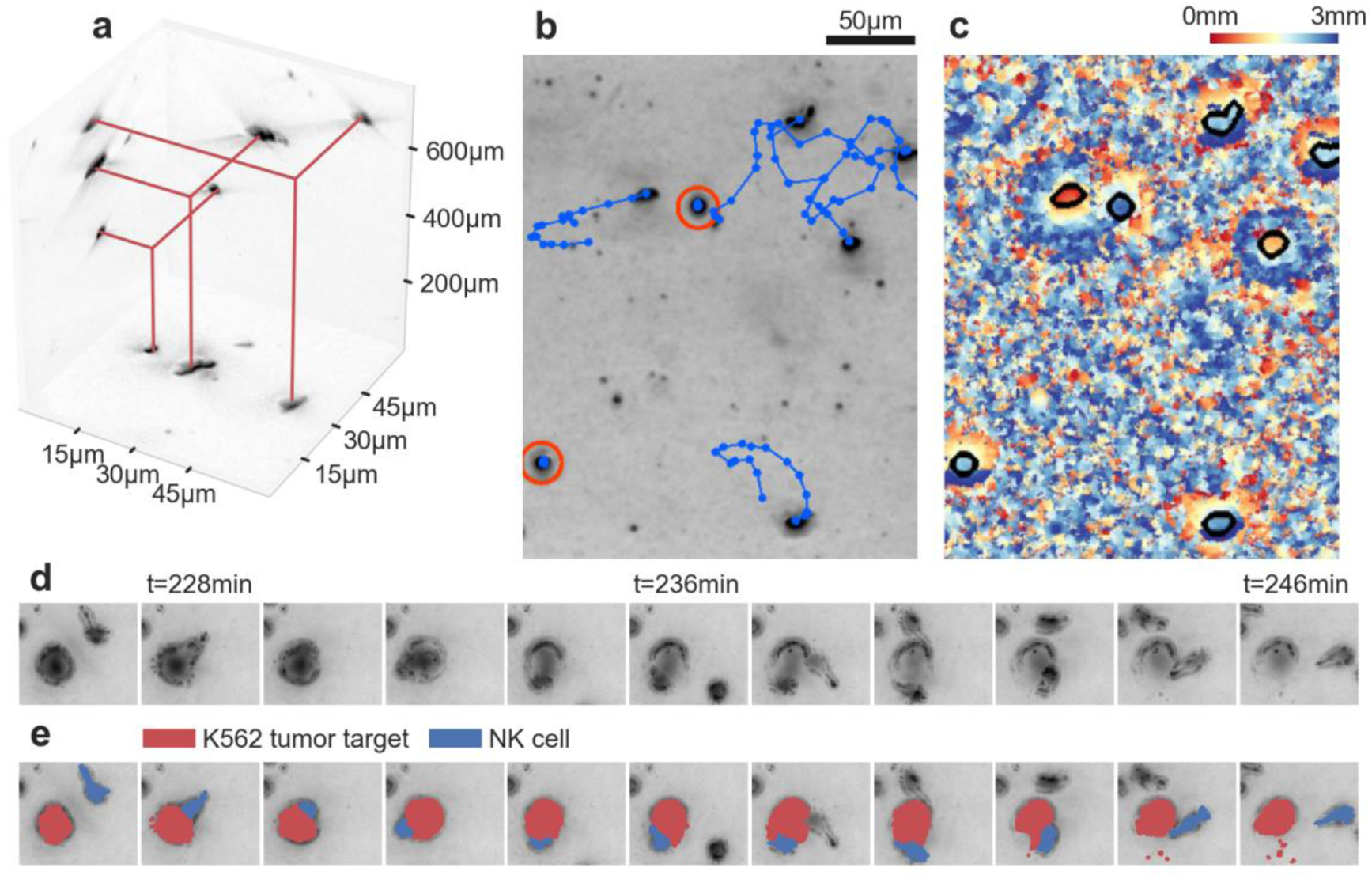
NK cell motility and cytotoxicity in a 3-D collagen gel. **a:** Minimum intensity projections along the x/y/z axes of a bright-field image stack of NK cells migrating through a 3-D collagen gel. In the projections, the NK cells appear as dark spots. **b:** Exemplary minimum intensity projection along the **z**-axis. Cell trajectories over 30 min are indicated in blue, non-motile cells are marked red. **c:** False-color representation of the **z**-position of every pixel in the minimum projection. This information is used to estimate the **z**-position of motile NK cells (cell outlines are indicated in black). **d:** Time-lapse image sequence of an NK cell-mediated killing of a K562 tumor cell. **e:** Same as (d), with colored cell morphologies as a guide for the eye.

For analysis, we detect individual NK cells using a convolutional neural network that is trained on 70 manually labeled minimum/maximum intensity projections with approximately 100 cells in each projection. We assume a minimum diameter of non-motile NK cells of 4 µm to separate them from smaller cell fragments (Supplementary Figure 2, Supplementary Video 4). In addition, we perform data augmentation by image flipping and zooming. The network is based on the U-Net architecture (19). The labeling accuracy of the network is 94% (F1-score), which is comparable to the accuracy of trained humans. We then connect the x,y positions of detected NK cells between subsequent images to obtain migration trajectories (Fig. 1b, Supplementary Video 1,2). An NK cell is classified as motile if it moves away from its starting point by 13 µm or more within the 5 min measurement time. The cell speed is determined as the diagonal of the bounding box of each cell trajectory divided by the measurement time of 5 min. Directional persistence is determined as the average cosine of the turning angles between consecutive cell movements. Zero persistence corresponds to random motion, whereas a persistence of unity corresponds to ballistic motion.

As the expanded cell populations contain small fractions of other cell types (as measured with flow cytometry, Fig. 2b), we perform confocal microscopy of an expanded cell population that is stained using CD56-APC and CD3-Alexa488 antibodies and is embedded in a 3-D collagen gel. We find that 82% of all motile cells are NK cells (Supplementary Figure 1).

**Figure 2:**
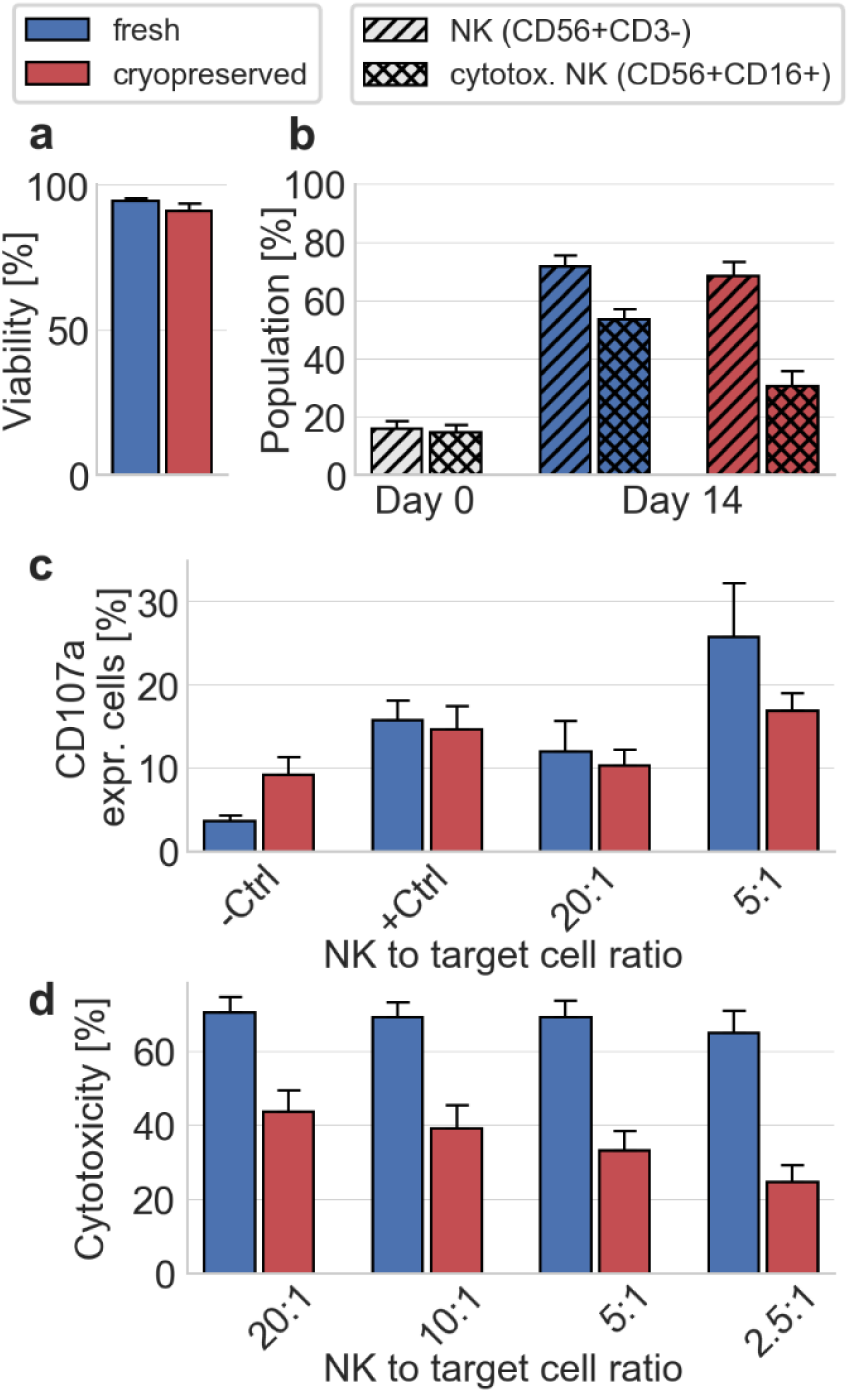
NK cell expansion and NK cell function in standard assays. **a:** NK cell viability of fresh and cryopreserved NK cells as measured by flow cytometry (p=0.36; n=9). **b:** Fraction of NK cells (single-hatched) and cytotoxic NK cells (double-hatched) within PBMCs before expansion (Day 0; white bars), and after expansion (Day 14) for fresh (blue) and cryopreserved (red) samples (n=10). The increase in the fraction of NK cells and cytotoxic NK cell after expansion is significant (p=0.002 for both conditions). Cryopreservation does not decrease the fraction of NK cells (p=0.49) but decreases the fraction of cytotoxic NK cells (p=0.004). **c:** Percentage of CD107a-expressing NK cells for different NK to target cell ratios, as measured in a degranulation assay (n=8). Control measurements are conducted in the absence of target cells. PMA/Iono is added for positive control experiments. Differences in the degranulation between fresh and cryopreserved NK cells do not reach statistical significance (-Ctrl: p=0.055, +Ctrl: p=0.74, 20:1: p=0.74, 5:1: p=0.38). **d:** NK cell cytotoxicity as measured in a chromium release assay for different NK to target cell ratios (n=10). Differences between fresh to cryopreserved NK cells reach statistical significance for all conditions (20:1: p=0.014, 10:1: p=0.006, 5:1: p=0.004, 2.5:1: p=0.002). For fresh NK cells, there is no significant correlation between NK-to-target cell ratio and cytotoxicity (correlation coefficient *ρ*=0.14, p=0.38), but for cryopreserved NK cells, cytotoxicity decreases for lower NK-to-target cell ratios (correlation coefficient *ρ*=0.41, p=0.009).

### 3-D cytotoxicity assay

We mix 600,000 fresh or cryopreserved NK cells together with 120,000 K562 cells in 2.5 ml of collagen (1.2 mg/ml) as described for the 3-D motility assay, except that z-stack time-lapse imaging is performed at four positions in each well in parallel, at a rate of 2 frames per min for 15 hours in total.

To automatically detect the number of motile NK cells as well as the number of live and dead K562 cells in each frame, we train a convolutional neural network with 48 minimum and maximum intensity projection images in which all motile and non-motile NK cells as well as live and dead K562 cells are manually labelled. Living K562 cells are characterized by a round appearance and small (1-2 µm) wiggling motion. When a K562 cell, after being contacted by an NK cell, starts to bleb, change shape, shrink, expand, stop wiggling, or change its bright-field contrast due to refractive index changes, we consider the cell as dead or as undergoing lysis or apoptosis (Fig. 1d,e; Supplementary Video 3). We complement this manually labeled data set with 11 images in which K562 cells are fluorescently labeled using Hoechst 33342 dye (incubation with 2.5 µg/ml for 20 min followed by 2 washing steps in PBS), 11 images in which NK cells are fluorescently labeled using Hoechst 33342 dye, and one image of dead K562 cells in which cell death is induced by adding 0.01 % TritonX. The complete data set is used to train a convolutional neural network based on the U-Net architecture (19). As non-motile NK cells are smaller but otherwise appear similar to K562 cells, we only consider K562 cells that are larger than 14 µm in diameter for the analysis (Supplementary Figure 2). An example demonstrating the performance of the automated detection and labelling is shown in Supplementary Video 4. For evaluating NK cytotoxicity, each NK cell-mediated killing event identified by the network is visually inspected and confirmed using the image-annotation software ClickPoints (20).

To evaluate the accuracy of determining target cell death in brightfield images, we add the dye NucRed Dead 647 (Ready Probes, Thermo Fisher) to the media immediately after the polymerization of the collagen gel and also at the end of the measurement (Supplementary Figure 3). At the beginning and at the end of the 15 h measurement period, an image stack is recorded. All K562 cells that are classified as living based on the bright field criteria are stained negative. Therefore, the false negative error rate is 0% (none of the dead cells are labelled falsely as living). However, from the K562 cells which are classified as dead based on the bright field criteria listed above, 10.6% are stained negative, possibly because they still have an intact cell membrane despite undergoing apoptosis. Therefore, we potentially overestimate the killing rate (see below) by up to 10%.

We describe the decline in the concentration of live K562 cells due to NK cell-mediated killing during the 15 h observation time as a first-order enzyme reaction of the form [*NK*] + [*K*562_*live*_] →^*k*^ [*NK*] + [*K*562_*dead*_], whereby the NK cells serve as the enzyme catalyzing this reaction, and *k* is the reaction rate (the killing efficiency). We assume that *k* remains constant and that the cytotoxicity of NK cells does not exhaust over time, which is an oversimplification but justified by the high ratio of NK cells to target cells. Moreover, we consider the average concentration of motile NK cells ([*NK*]) as being constant throughout the measurement. Finally, we assume that the reverse reaction does not take place, implying that K562 cells do not recover once they are undergoing apoptosis or lysis. Therefore, if a K562 cell forms small temporary blebs that disappear after a while so that the cell appears alive for the remainder of the experiment, this cell is counted as living.

The dynamics of the above killing reaction can be described with a first-order differential equation, which has the solution 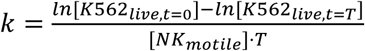. The only free parameter is the killing efficiency *k* that we compute from the concentration (# of cells per (100µm)^3^ of gel volume) of live K562 cells at the beginning (t=0) and the end of the experiment (t=T), and the average concentration of motile NK cells. As individual experiments *i* have different cell concentrations, we compute weighted mean killing efficiencies 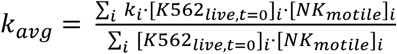 for both fresh and cryopreserved NK cells. Below, we also report an apparent or composite killing rate for all NK cells regardless of their ability to move, which is computed according to 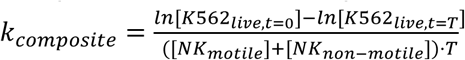. Since only motile NK cells can reach their target cell to kill, this composite killing rate combines cytotoxicity and motility in one parameter.

### Statistical tests

Unless noted otherwise, all statistical tests use the two-sided non-parametric Wilcoxon-signed rank test for paired data. The correlation between NK-to-target ratio and cytotoxicity reported in Fig. 2d is determined by Spearman’s Rank Order Correlation; the corresponding test statistic is based on a t-distribution to account for the small sample size and to avoid over-rejection. Significant differences for the killing efficiency reported in Fig. 4d are determined by bootstrapping (two-sided test). Differences are considered as statistically significant for p<0.05.

## Results

Before investigating NK cell function after cryopreservation, we confirm by live-dead staining and flow cytometry that cell viability is not affected by the process of freezing and thawing (Fig. 2a). The 14-day expansion protocol significantly increases the fraction of NK cells (identified as CD56+ CD3-) within the PBMC population by a factor of 4, which remains unchanged after cryopreservation (Fig. 2b). However, cryopreservation results in a significant decrease of the CD16+ subpopulation of NK cells (Fig. 2b). This subpopulation is also referred to as activated or cytotoxic NK cells as the CD16 surface receptor is a known mediator of NK cell cytotoxicity.

To assess whether the relative decrease in the CD16+ subpopulation of NK cells affects the potential antitumor activity of the whole expanded NK cell population, we perform a CD107a degranulation assay. This assay tests the ability of NK cells to fuse lytic granules with their membrane, which is an essential step to induce target cell death. We find no significant difference in the expression of CD107a between fresh and cryopreserved NK cells, indicating that cryopreserved NK cells retain the ability to induce target cell death (Fig. 2c).

Next, we evaluate target cell death directly in a co-culture of NK cells and K562 target cells across a range of NK-to-target cell ratios using a chromium-release cytotoxicity assay. We find a statistically significant (p<0.05) decrease in target cell death after cryopreservation compared to fresh NK cells (Fig. 2d). This decrease in cytotoxicity may result from two different mechanisms: First, cryopreservation may affect NK cell activation, as indicated by the reduced number of CD16+ cells. Second, cryopreservation may reduce NK cell motility such that they cannot come in contact with target cells.

In support of the second mechanism, we find that the decrease in cytotoxicity after cryopreservation is more pronounced for smaller NK-to-target cell ratios (for the same number of K562 cells). Assuming that only a fraction of NK cells mediates the majority of target cell deaths (21), those NK cells would need to kill a larger number of K562 cells to retain the same overall cytotoxicity in cell populations that contain fewer NK cells. At the same time, those NK cells would need to migrate over larger distances to reach target cells. For fresh NK cells, by contrast, we find no statistically significant correlation between cytotoxicity and NK-to-target ratio (Fig. 2d), indicating that fresh NK cells can compensate for a decreasing NK-to-target cell ratio.

Since the chromium-release assay only reports the decreased cytotoxicity but not cannot measure NK cell motility directly, we perform a motility assay that assesses the fraction of an NK cell population that is motile when embedded in 3-D reconstituted collagen gels. We find that NK cell motility after cryopreservation in a 3-D environment is dramatically impaired after cryopreservation. Specifically, we find that 29.2% of fresh NK cells are motile, in line with reported findings for 2-D migration (22), while only 4.9% of cryopreserved NK cells are motile, corresponding to a 6-fold decrease in motility (Fig. 3a).

**Figure 3:**
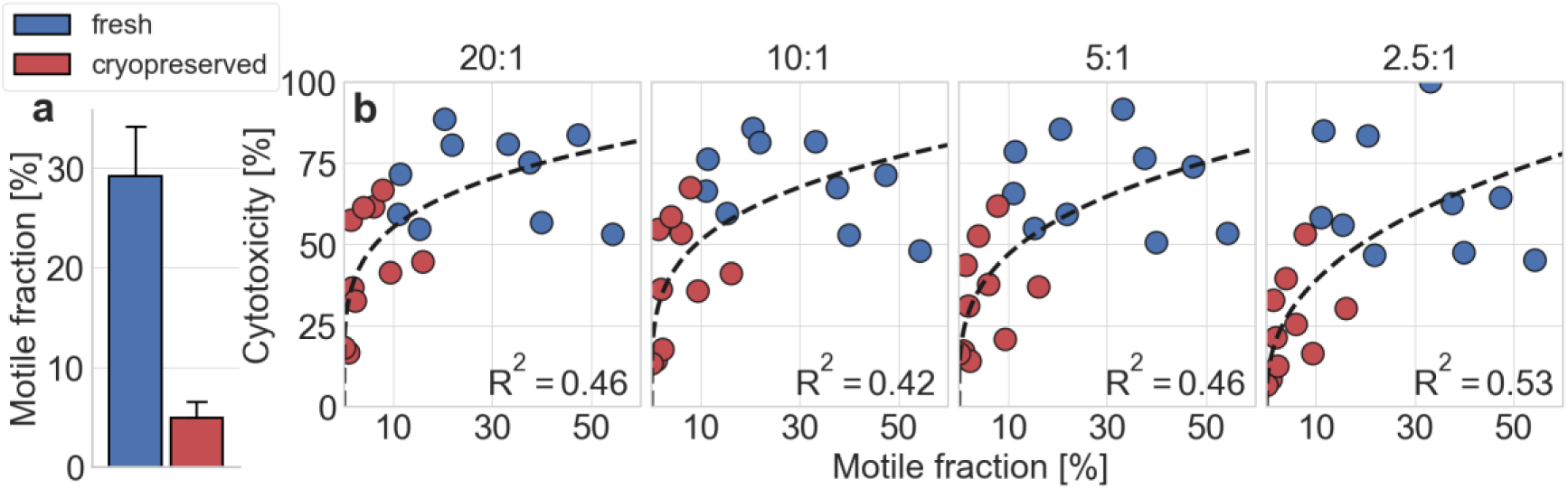
Influence of 3-D NK cell motility on cytotoxicity. **a:** Motile fraction of fresh and cryopreserved NK cells in a 1.2 mg/ml collagen gel (p=0.002; n=10). **b:** NK cell cytotoxicity (as measured in a chromium release assay) as a function of the motile NK cell fraction for fresh (blue) and cryopreserved (red) NK cells. Different NK-to-target cell ratios are noted above each graph. Dashed lines indicate a power-law fit of the form *f*(*x*) = *a* ⋅ *x*^*b*^. *R*^2^values are computed in log-log-space. Statistical significance assuming a power-law exponent of *b* = 0 as Null hypothesis: 20:1: p=0.0013; 10:1: p=0.0028; 5:1: p=0.0013; 2.5:1: p=0.00045.

As chromium-release assays and motility assays are performed pair-wise, we can quantify the relationship between cytotoxicity and motile cell fraction in 3-D collagen gels for individual samples (specified by donor and expansion). We find that cytotoxicity increases with motile fraction according to a power-law relationship with exponents ranging from 0.2-0.4. This relationship reaches statistical significance, with a coefficient of determination of *R*^2^ ≥ 0.42 for all NK-to-target cell ratios between 20:1 and 2.5:1 (Fig. 3b). This finding supports the notion that NK cells are able to kill target cells beyond their immediate neighbors in a cell pellet (e.g. in a chromium release assay), for which the ability to migrate is of advantage.

We next investigate whether the small remaining population of motile NK cells after cryopreservation retains full cell function. First, we characterize cell speed and directional persistence of fresh and cryopreserved motile NK cells within 3-D collagen gels and find no significant difference (Fig. 4a,b). Thus, NK cells that remain motile after cryopreservation retain their normal exploration behavior within tissue.

**Figure 4:**
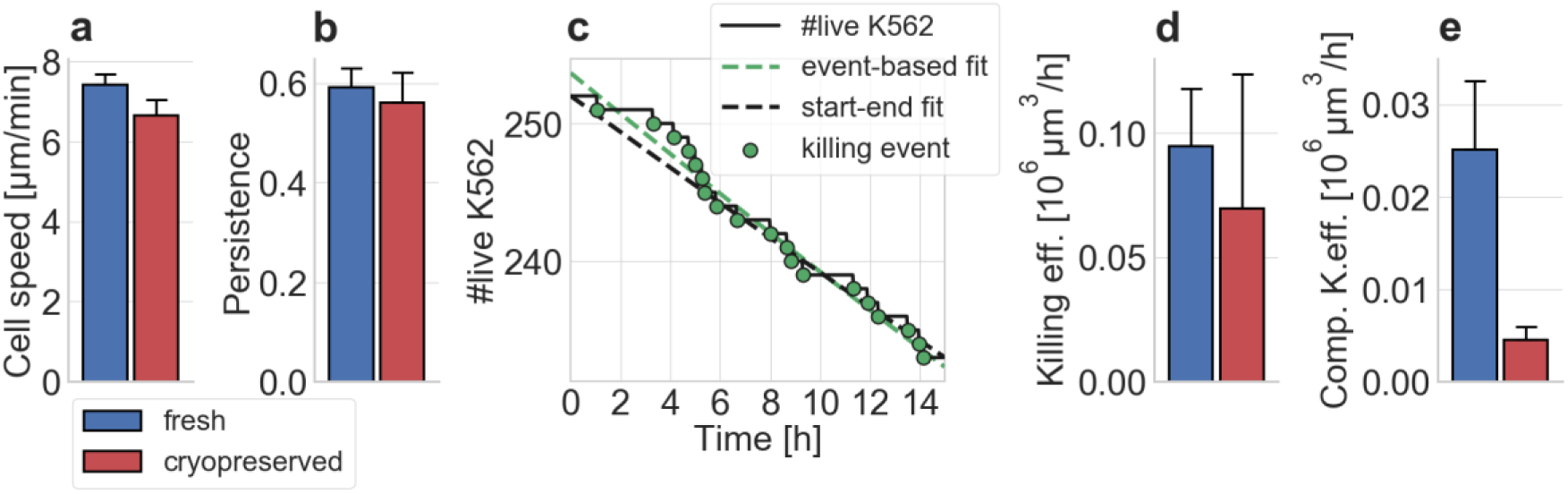
NK cell migration and cytotoxicity in 3-D collagen gels. **a:** Cell speed (mean ± se) (measured in the x-y-imaging plane) of the motile fraction of fresh and cryopreserved NK cells embedded in a 3-D collagen gel (p=0.076; 1248 fresh and 122 cryopreserved motile NK cells from n=10 independent experiments). **b:** Mean directional persistence of the motile fraction of fresh and cryopreserved NK cells embedded in a 3-D collagen gel (p=1.0; n=10). **c:** Number of live K562 target cells (black line) in a representative experiment as a function of measurement time. Killing events are marked by green circles. The dashed green line indicates an exponential fit of the individual killing events, the black dashed line indicates an approximation using an exponential curve based only on the number of live targets at the beginning and the end of the measurement time. **d:** Estimated killing efficiency of fresh and cryopreserved motile NK cells against K562 target cells embedded in a 3-D collagen gel (p=0.56; n=6). **e:** Estimated composite killing efficiency of fresh and cryopreserved NK cells (considering both motile and non-motile NK cells) against K562 target cells embedded in a 3-D collagen gel (p=0.025; n=6).

Second, we evaluate the cytotoxic function of the motile NK cell fraction in 3-D collagen gels by identifying individual K562 target cell killing events in time-lapse image series (Supplementary Videos 3-5). The number of live target cells decreases exponentially over time, which can be modeled by a first-order differential equation (see Methods). The characteristic time constant of the exponential decay, which reports the killing rate and hence the cytotoxicity of the motile NK cells, can be estimated from the number of live target cells at the beginning and the end of the experiment (Fig. 4c). Normalizing this exponential decay rate to one hour as well as to one motile NK cell and one living target cell per 10^6^ µm^3^ volume of tissue (corresponding to a cube of 100 µm length), we arrive at a measure of killing efficiency that is not biased by varying cell concentrations or fractions of motile cells (Supplementary Figure 4). We find a 26% decrease of cytotoxicity in a 3-D environment after cryopreservation, but this difference is smaller than the difference detected in the classic cytotoxicity assays and is not statistically significant (Fig. 4d). Therefore, our data demonstrate that NK cells that remain motile after cryopreservation retain their cytotoxic cell function. However, if we compute a composite killing efficiency that considers both motile and non-motile NK cells, we find a significant decrease of cytotoxicity in a 3-D environment by a factor of 5.6 (Fig. 4e), which directly reflects the 6-fold decrease in NK cell motility.

## Discussion

In this study, we report that ex-vivo expanded NK cells retain their ability to induce target cell death after cryopreservation (as measured in a degranulation assay), but suffer a significant decrease of their cytotoxic function (as measured in a chromium-release assay). By complementing standard assays of NK cell function with measurements of NK cell motility and cytotoxicity on a single-cell level in a 3-D environment, we are able to attribute this decrease of cytotoxic function to an impairment of NK cell motility. Specifically, we find a dramatic 6-fold decrease in the fraction of motile NK cells after cryopreservation. We further show that the small remaining population of motile NK cells retains its cytotoxicity after cryopreservation.

These findings point to the cryopreservation step as a major cause for the current failure of NK cell therapy in patients with solid tumors, and that the fraction of motile NK cells is a crucial parameter when deciding on the number of cells to be administered to the patient. Finally, this work demonstrates that time-lapse imaging of NK cells and tumor cell killing events in a 3-D matrix may serve as a better predictor of NK cell function compared to standard cytotoxicity assays.

## Acknowledgements

We thank Dr. J.J. Bosch and Prof. D. Campana for providing the tumor cell lines.

## Author Contributions

C.V., B.F., and G.S. designed the study. S.R. performed cell expansion, cryopreservation, cytometry, degranulation and chromium-release assays. C.M., T.C., A.M., F.H., and N.B. developed and performed the 3-D motility and cytotoxicity assays. L.H., A.W., S.S., and S.R. developed the neural network-based analysis method. C.M., T.C., L.H., and S.S. performed data analysis. C.M. and T.C. generated the figures. C.M., B.F., and C.V. wrote the manuscript.

