## Supplementary Information for "Cryopreservation impairs cytotoxicity and migration of NK cells in 3-D tissue: Implications for cancer immunotherapy"

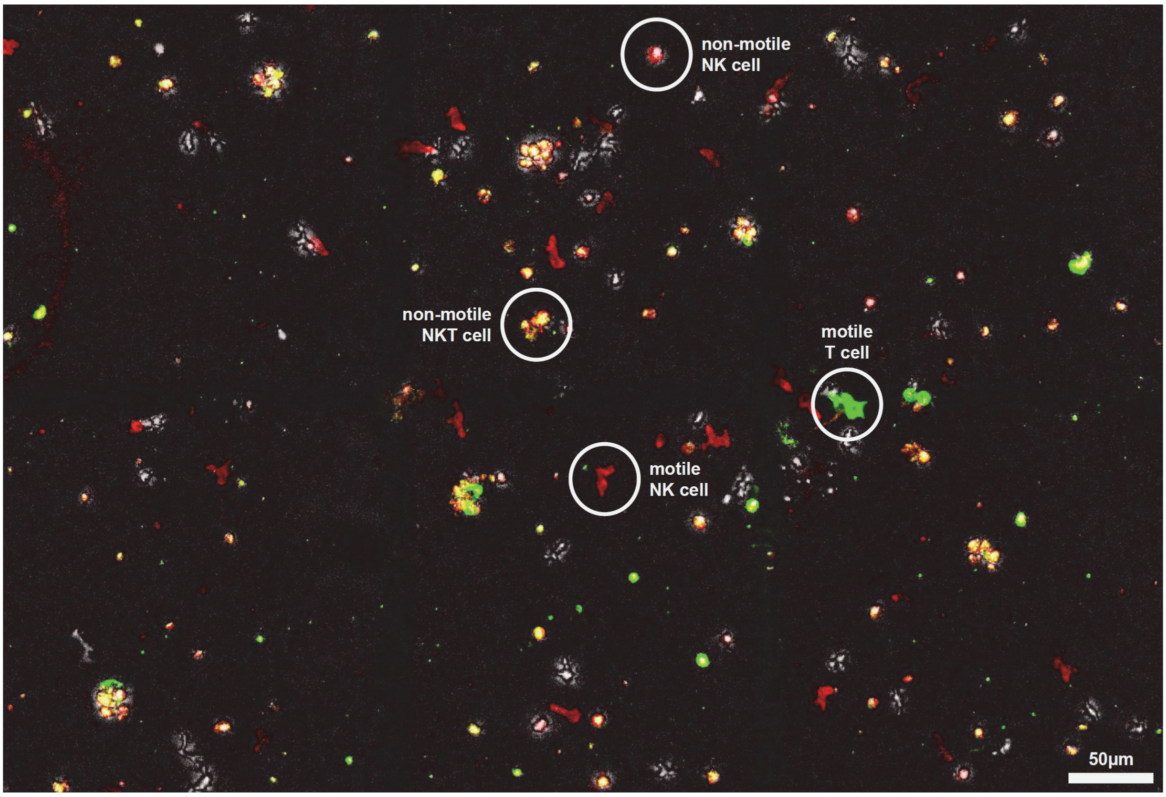


**Supplementary Figure 1:** Composite confocal maximum intensity projections of a cell population in a 3D collagen gel expanded from PBMCs. The brightfield maximum intensity projection at the beginning of the experiment is shown in grey-scale. The red overlay shows the maximum intensity projection of a CD56-APC staining that was recorded 10 minutes after recording the brightfield image stack. The green overlay shows the maximum intensity projection of a CD3-Alexa488 staining that was also recorded 10 minutes after recording the brightfield image stack. Thus, cells that appear red on a dark background are motile NK cells, while cells that appear red on a bright background have not moved in 10 minutes and represent non-motile NK cells. Likewise, cells that appear yellow (double positive for CD56 and CD3) are NKT cells, and cells that appear green are T cells. Based on three field-of-views, we find that 82% of all motile cells are NK cells.

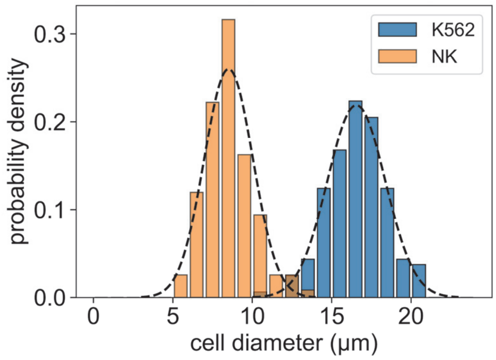


**Supplementary Figure 2:** Size distribution of K562 cells and non-motile (round) NK cells (K562 n = 162, NK n = 117). The overlap in the size distributions is less than 2%.

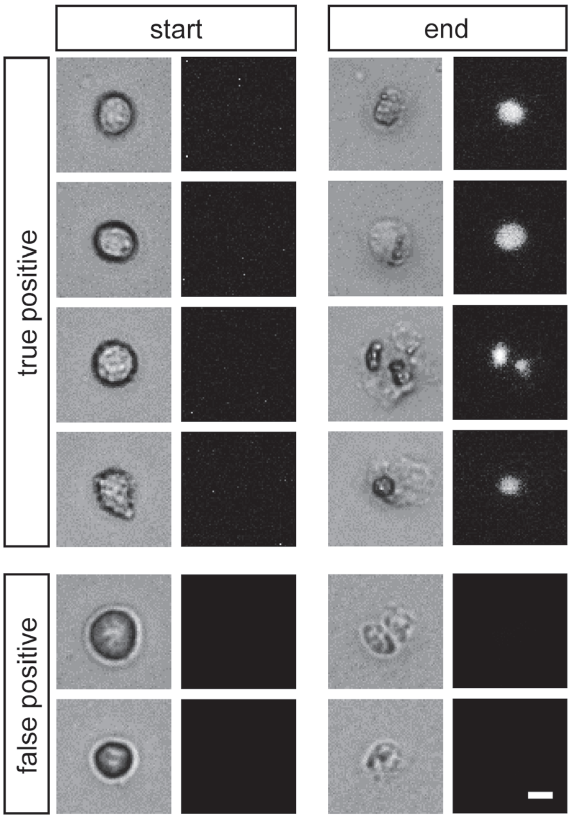


**Supplementary Figure 3:** Killing events as evaluated based on bright field images are verified with NucRed Dead 647 staining (Ready Probes, Thermo Fisher). 89.4% of all K562 cells that are classified as “dead” based on bright field criteria are stained positive (true positives). 10.6% of all K562 cells that are classified as “dead” based on bright field criteria are stained negative (false positives). These cells may possibly be undergoing apoptosis but may still have an intact cell membrane. In n=196 evaluated cells, we found no false negative events (no cell that was classified as “living” stained positive). Scale bar: 10 μm.

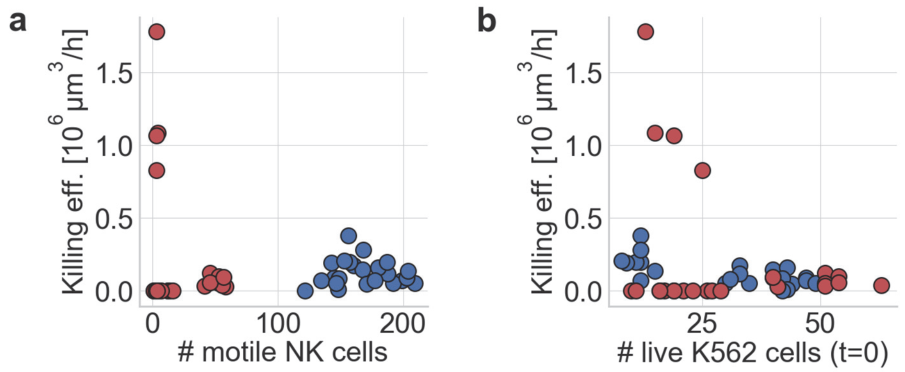


**Supplementary Figure 4:** **a:** Killing rate for individual fields-of-view as a function of the number of motile NK cells as determined by the 3-D cytotoxicity assay. **b:** Killing rate for individual fields-of-view as a function of the number of K562 target cells. We find no systematic dependence of the killing rate on either the number of NK cells or the number of target cells, demonstrating that the killing rate estimation is not biased by different concentrations of NK cells and target cells.

**Supplementary Video 1:** Series of minimum intensity projections through a 3-D collagen gel with embedded, freshly expanded NK cells. The video spans a measurement time of 5 minutes. Automatically evaluated cell trajectories are color-coded.

**Supplementary Video 2:** Series of minimum intensity projections through a 3-D collagen gel with embedded, cryopreserved expanded NK cells. The video spans a measurement time of 5 minutes. Automatically evaluated cell trajectories are color-coded.

**Supplementary Video 3:** Series of minimum intensity projections through a 3-D collagen gel with embedded, cryopreserved expanded NK cells and K562 target cells. Video shows a cropped region around an NK cell-mediated killing event of a K562 target cell (approx. 10 h after the beginning of the measurement) and spans the complete 15 h of measurement time.

**Supplementary Video 4:** Same as in Supplementary Video 3, but with automatic labeling using a convolutional neural network. Blue regions indicate live K562 cells, red regions indicate motile NK cells, green regions indicate non-motile NK cells as well as smaller cell fragments, and magenta regions indicate a contact between NK cell and target cell.

**Supplementary Video 5:** Same as in Supplementary Video 3, but showing the full field-of-view.
